# FK506-binding protein FklB is involved in biofilm formation through its peptidyl-prolyl isomerase activity

**DOI:** 10.1101/2020.02.01.930347

**Authors:** Chrysoula Zografou, Maria Dimou, Panagiotis Katinakis

## Abstract

FklB is a member of the FK506-binding proteins (FKBPs), a family that consists of five genes in *Escherichia coli*. Little is known about the physiological and functional role of FklB in bacterial movement. In the present study, FklB knock-out mutant Δ*fklB* presented an increased swarming and swimming motility and biofilm formation phenotype, suggesting that FklB is a negative regulator of these cellular processes. Complementation with Peptidyl-prolyl isomerase (PPIase)-deficient *fklB* gene *(Y181A*) revealed that the defects in biofilm formation were not restored by *Y181A*, indicating that PPIase activity of FklB is modulating biofilm formation in *E. coli*. The mean cell length of Δ*fklB* swarming cells was significantly smaller as compared to the wild-type BW25113. Furthermore, the mean cell length of swarming and swimming wild-type and Δ*fklB* cells overexpressing *fklB* or *Y181A* was considerably larger, suggesting that PPIase activity of FklB plays a role in cell elongation and/or cell division. A multi-copy suppression assay demonstrated that defects in motility and biofilm phenotype were compensated by overexpressing sets of PPIase-encoding genes. Taken together, our data represent the first report demonstrating the involvement of FklB in cellular functions of *E. coli*.

## Introduction

The previously prevalent view of bacteria development was considered to be the planktonic form of life. That is, unicellular organisms that grow as individual entities. This view has changed as it has been found that bacteria, under given conditions, behave as multicellular groups that grow on nutrient-rich surfaces, secrete a polysaccharide material, through a process called swarming motility. Swarming motility is mainly driven by rotating flagella, and swarming bacteria generally appear to be elongated as a result of cell division suppression [1][2]. In contrast to swarming, swimming describes a mode in which cells move within aqueous environments, not in groups, but independently, by operating their rotating flagella [3]. A comparable type of multicellular behavior is the biofilm formation where bacteria form sessile communities and disperse by secreting proteins and surfactants extracellularly [4]. Biofilms are complex systems and can be composed of multiple species [5]. Environmental conditions and coordinated life cycles can affect or set off heterogeneity and include, among other, expression of genes and proteins, as well as post-translational protein modifications (PTMs), that could alter environmental sensing and signal transduction [6][7]. Various PTMs such as glycosylation, N-terminal modifications and phosphorylation are few of the functional properties of Peptidyl-prolyl *cis*/*trans* isomerases (PPIases). PPIases, being ubiquitous among all organisms, are key regulators of numerous highly important biological processes; they accelerate the rate of *in vitro* protein folding and they have the ability to bind proteins and act as chaperones.

Additionally, PPIases catalyze the folding of newly synthesized protein targets, particularly those that have peptide bonds in the *trans* conformation. They are also able to alter the structure and conformation of mature proteins thus affecting their intermolecular interactions [8]. In bacteria and other organisms, there are three characterized PPIase subfamilies; the Cyclophilins, the FK506-binding proteins (FKBPs), and the Parvulins. *E. coli* FKBP family consists of five genes; *fkpA, fkpB, fklB, slyD* and *tig*, none of which is essential for growth [9]. FKBPs are found to be involved in a diverse series of cellular processes such as cell division [10], stress response regulation and development [11], gene regulation through transcription and translation [12], and most importantly, virulence and pathogenicity [13].

*E. coli* FklB (or FKPB22) possesses PPIase activity, exists in solution as a homodimer and shares a significant homology with the protein Mip (macrophage infectivity potentiator) that is identified in a number of human pathogenic bacteria, such as *Legionella pneumophila, Neisseria gonorrheae*, and *Chlamydia trachomatis* in the psychrotrophic bacterium *Shewanella* sp. SIB1 [14] but also in the plant pathogen *Xanthomonas campestris* [15]. *Shewanella* SIB1 FKBP22 is composed of two monomers that are connected at their N-termini, bearing a V-shaped structure. Within the monomer, a 40-residue long a-helix separates the N- and C-terminal domain [16]. An almost identical tertiary structure appears to be assumed by the *E. coli* FklB [17]. Data suggest that the probable PPIase binding site of SIB1 FKBP22 for a protein substrate is located at its C-terminal domain. Abrogation of SIB1 FKBP22 PPIase activity did not significantly affect its chaperone function [18].

This paper describes a new approach to investigate the physiological role and assess the PPIase and chaperone function of FklB through a series of phenotypic methods. We focused on genetic and biochemical approaches to assess the swarming and swimming motility, biofilm and cell length phenotypes in *E. coli*, caused either by the loss of *fklB* or by the overexpression of *fklB* and of PPIase-deficient *fklB* (*Y181A*) gene. We found that deletion of *fklB* resulted in an enhancement of motility and biofilm, as well as a decrease of swarming cells’ length. Complementation with *fklB* gene, in the mutant strain ΔFklB, suppressed the *fklB* deletion motility and biofilm phenotype, while overexpression of *Y181A* suppressed only the motility phenotype. Overexpression of *fklB or Y181A* gene exhibited opposite effects on the mean cell length of swarming and swimming cells. We also used a multi-copy suppression approach to assess if overexpression of other PPIase-encoding genes may suppress the Δ*fklB* strain motility and biofilm phenotypes.

## Results & Discussion

### PPIase and chaperone activity of FklB

Initially, we examined the PPIase activity of FklB with a standard PPIase assay of isomer-specific proteolysis by chymotrypsin, described by Kofron [19]. Based on the protein alignment of *E. coli* FklB (Ec_4207) with the fully characterized human FkpB12 (hfkpB12), we located the FklB’s putative active sites. We then constructed an active site mutated form of FklB, Y181A, which we used in the PPIase assay in comparison to the wild-type FklB. N-succinyl-Ala-Ala-Pro-Phe-pnitroanilide was used as a substrate known to mimic the internal peptidyl-prolyl moiety of proteins containing proline.

We found that Y181A had no measurable isomerase function, whereas the catalytic efficiency of FklB was 1.50 ± 0.0013 (Kcat/Km), suggesting that substitution of Y181 had a significant effect on its activity (Fig. 1A). This result suggests that Y181A, located on its C-terminal domain, is involved in the catalytic function of the enzyme. Previous findings have demonstrated that the catalytic efficiencies of other mutant forms of FklB indicate that W157 and F197 are also critically important for the isomerase activity of SIB1 FKBP22 [20].

**Figure 1.**
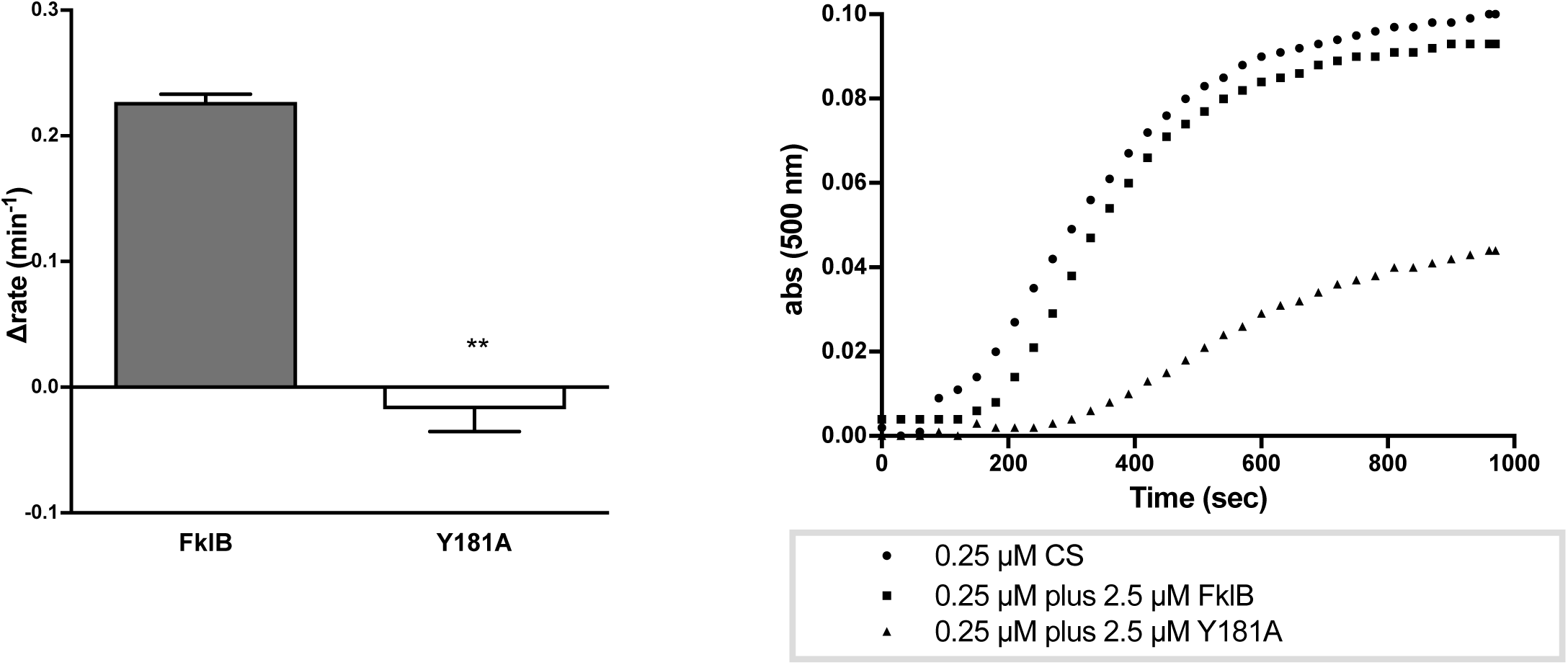
Prolyl isomerase and chaperone activity is lost in mutant Y181A compared to the wild-type protein FklB. **A.** PPIase activity of 0.25 uM FklB and Y181A. Mean values were obtained from three independent replicates and error bars represent standard errors. **B.** Thermal aggregation of citrate synthase in the absence (•) and presence of (▪) 0.25 uM FklB or 0.25 uM Y181A (▲). The results are representative of three series of measurements carried out with different preparations of enzymes.

Next, we sought to determine the chaperone activity of FklB by measuring its ability to suppress the thermal aggregation of citrate synthase (CS) [21], while also evaluating any effects on it, caused by the induced active site mutation. CS tends to aggregate at high temperatures because of hydrophobic interactions between unfolding intermediates and results in the formation of high molecular weight particles. Proteins with a chaperone function are able to recognize and bind to these unfolding intermediates and therefore keep their concentration low in solution. We observed that wild-type FklB presents a chaperone function, whereas the inhibition of the CS aggregation from Y181A appears to increase in proportion to its concentration (Fig. 1B).

However, we noticed an increased ability of Y181A to inhibit the formation of CS aggregates, in comparison to the wild-type FklB. This may be an indication that the loss of FklB’s PPIase activity improves its function as a chaperone.

It has been previously shown that the PPIase and chaperone activity of SIB1 FKBP22 reside in two structurally unrelated domains, but not necessarily functionally independent domains. Mutations at its PPIase active site do not critically affect its chaperone function, an indication that SIB1 FKBP22 does not require PPIase activity for protein folding. However, the authors highlight the importance and requirement of the chaperone domain for the PPIase activity, as a way of enabling the formation of folding intermediates [20]. Several other studies have also indicated that the presence of chaperone activity improves PPIase activity [22] [23]. The substrates are bound to the chaperone site and are subsequently transferred to the PPIase site, where the peptidyl-prolyl bonds of the proteins are being isomerized. Protein molecules, that exited the PPIase site with an incorrect peptide-prolyl bond, are re-attached to the chaperone region and the procedure is repeated [23]. Therefore, it could be assumed that the chaperone activity of FklB may offset a high PPIase activity and concurrently the loss of PPIase activity might allow structural changes that increase the chaperone activity. Elucidating the structural relationship and association of the two functions is very important in order to uncover the role of the PPIase family in major cellular processes.

### Role of FklB in swarming and swimming motility

The role of FklB was examined under swarming and swimming conditions, by inoculating the center of swarming (LB-glucose, 0.5% agar) and swimming plates (LB, 0.3% agar) with liquid cultures expressing and/or lacking the *fklB* gene (Fig. 2). We found that the mutant ΔFklB strain formed considerably larger swarming and swimming colonies in comparison to the control strain, BW25113, indicating that the loss of FklB is responsible for the observed phenotype (Fig. 2A-B). In order to validate that FklB functions as a swarming and swimming motility repressor we examined the phenotypes of the ΔFklB strain and of the control strain overexpressing the *fklB* gene (strain BW25113(FklB)). We found that the ΔFklB(FklB) reverted the hyper-swarmer or hyper-swimmer phenotype to wild-type, while BW25113(FklB) further suppressed the phenotype beyond the wild-type levels (Fig. 2A-D). This observation supports our initial hypothesis that the lack of FklB was the causative factor of the increased swarming and swimming motility. Growth rates of ΔFklB or BW25113 (FklB) strain liquid cultures were comparable to the control’s, suggesting that the increased motility phenotype was not attributed to an increased growth.

**Figure 2.**
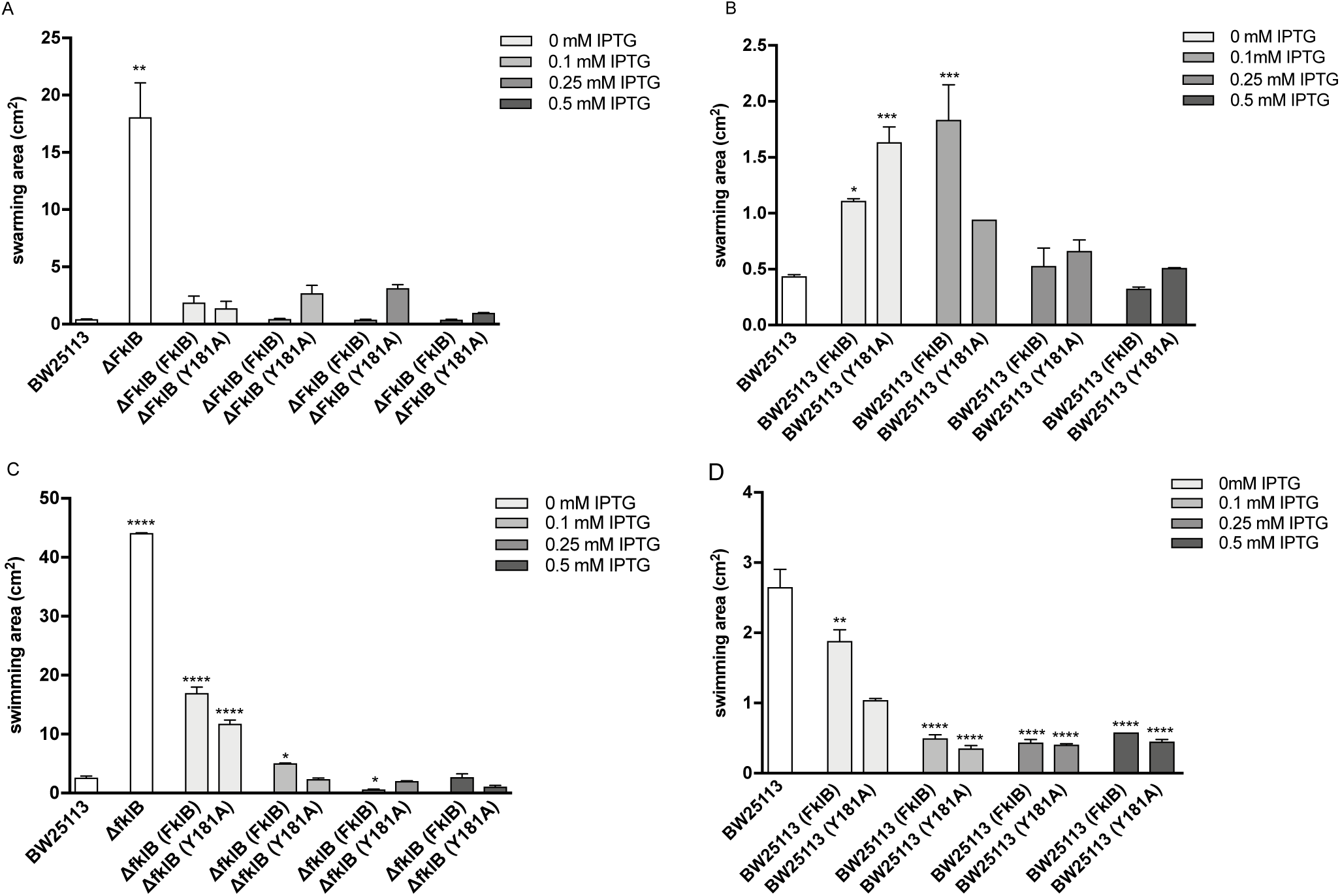
Prolyl isomerase activity of FklB causes a suppression of the swarming and swimming phenotype in *E. coli*. Swarming (A) and swimming (C) area of the ΔFklB mutant strain that overexpresses FklB and Y181A and swarming (B) and swimming area (D) of BW25113 that overexpresses FklB and Y181A, compared to the wild-type BW25113. Mean values were obtained from four independent replicates, and error bars represent standard errors. Statistical comparisons were made using ANOVA followed by Dunnett’s multiple-comparison test. Asterisks indicate statistically significant differences (P < 0.05).

Subsequently, we checked the involvement of the PPIase activity of FklB protein in swarming and swimming motility by following the same conditions. Strain ΔFklB(Y181A) did not seem to differ from the corresponding strain ΔfklB(FklB), even in highest IPTG concentrations, as both displayed no motility on swarm or swim plates, suggesting that the PPIase activity of FklB is not likely to be involved in the mechanism. Overall, we found that the expression of FklB and its mutant form, Y181A, is able to restore the wild-type phenotype at all IPTG concentrations (0.1-0.5 mM). This confirms that the presence of FklB is indispensable for maintaining a normal phenotype, perhaps through pre- and post-translational modifications or indirect target protein interactions that control a wide range of cellular processes, including motility [24] [25]. The negative regulation of swarming and swimming motility in *E. coli* by certain PPIase family members was previously shown [26][27]. FkpB proteins are found to be involved in bacterial motility, for example, an increase in the transcript levels of *fklB* gene was observed in *P. mirabilis* swarming cells [28]. Another example is the GldI protein of the microorganism *F. johnsoniae*, a lipoprotein homologous to FKBPs, essential for gliding mobility [29]. Although the structure of almost all FKBP proteins has been extensively studied, our knowledge about their biological role still remains limited. We already know that FKBPs catalyze the refolding of peptides preceding proline at polypeptide chains, as well as that all exhibit some PPIase activity, but there are still several unanswered questions about their physiological role.

### FklB is suppressing *E. coli*’s biofilm formation ability

Swarming motility and biofilm formation relationship seems to be complex and although both conditions share some common constituents, they greatly differ. Specifically, the use of flagella is necessary for biofilm initiation, but motility is also required for its initiation, as well as dispersion and release of bacteria [30]. However, it is not clear whether there is an inverse regulation of swarming motility and biofilm formation, as conflicting data have been published. For example, an increased EPS production suppressed swarming motility, but enhanced biofilm formation among laboratory isolates [31]. Biofilm formation, as well as swarming and swimming motility, was suppressed by overexpressing the cyclophilin PpiB. However, this involvement of PpiB in the biofilm formation phenotype does not involve its prolyl isomerase activity [32]. Biofilm formation was explored for FklB in order to elucidate the role of its PPIase function in this multicellular behavior, but also to investigate into its relation to swarming. To this means, we initially compared the biofilm formed by the control BW25113 and the mutant strain ΔFklB and we noticed that the ΔFklB strain was capable of a greatly increased biofilm formation (Fig. 3). We hypothesized that the increased biofilm formation by ΔFklB was attributed to the absence of the FklB protein and in order to clarify this we tested the biofilm formation under the same conditions of the strain ΔFklB(FklB) as well as the strain BW25113(FklB). Indeed, we noticed the restoration of the wild-type phenotype, when FklB was expressed at intermediate IPTG concentrations (0.1-0.25 mM), in the mutant strain (ΔFklB(FklB)) and in the wild-type strain (BW25113(FklB)). Interestingly, we further detected a biofilm repression phenotype when FklB was overexpressed (0.5 mM IPTG), either in BW25113(FklB) or in ΔFklB(FklB) strain, suggesting that FklB bears a key role in biofilm formation (Fig. 3A-B).

**Figure 3.**
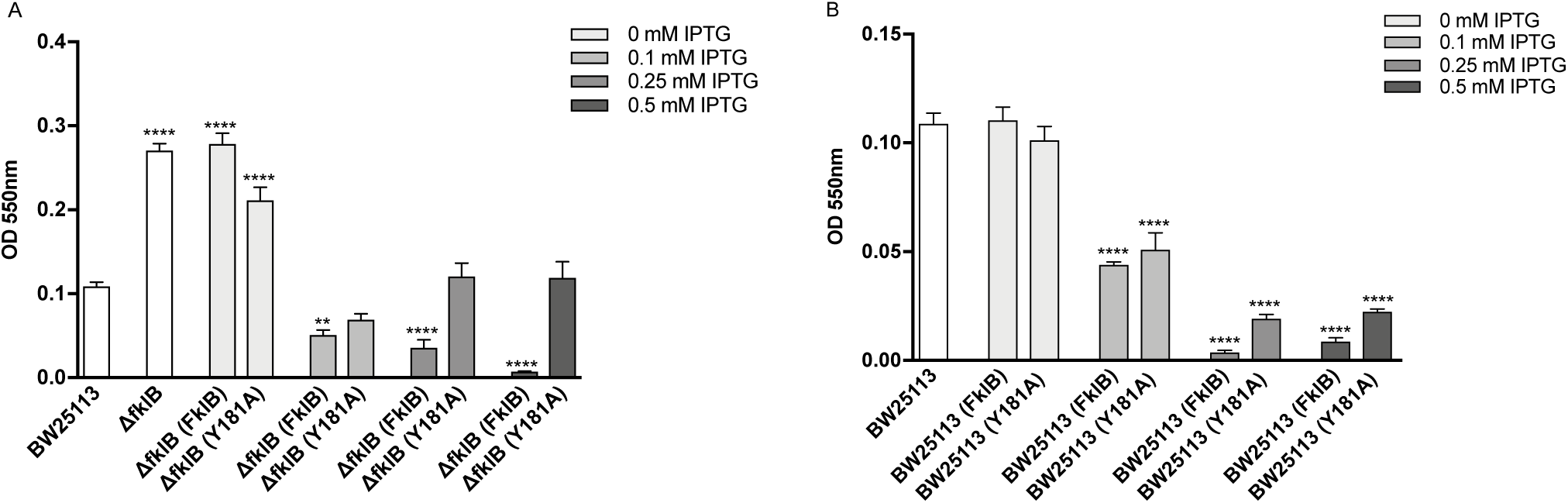
Prolyl isomerase activity of FklB causes a suppression of the biofilm phenotype in *E. coli*. Biofilm formation of the ΔFklB mutant strain that overexpresses FklB and Y181A (A) and of the BW25113 that overexpresses FklB and Y181A (B), compared to the wild-type BW25113. Mean values were obtained from four independent replicates, and error bars represent standard errors. Statistical comparisons were made using ANOVA followed by Dunnett’s multiple-comparison test. Asterisks indicate statistically significant differences (P < 0.05).

Similarly, we tested the strains that overexpress the mutant protein Y181A, ΔFklB(Y181A) and BW25113(Y181A). The results showed that the biofilm of strain ΔFklB(Y181A), did not differ from the mutant strain ΔFklB, even in the presence of high levels of IPTG (0.25 and 0.5 mM). The overexpression of the mutant Y181A did not cause a restoration of the wild-type phenotype. Based on these results, we can conclude that FklB’s PPIase activity is involved in this multicellular behavior (Fig. 3A).

Additionally, the strain BW25113(Y181A), in the absence or presence of low levels of IPTG (0.1 mM), showed similar biofilm formation ability to the control BW25113. However, we noticed that even though the strain BW25113(Y181A) at 0.25 and 0.5 mM IPTG, showed an important decrease in biofilm formation, that decrease was slightly lower than BW25113 (FklB) ((Fig. 3B). These observations seem to suggest that FklB has an important role in suppressing the biofilm formation phenotype of *E. coli* and that its PPIase activity is indispensable for this involvement.

### PPIase family members can functionally replace FklB in swarming, swimming and biofilm cells

Previous research has demonstrated that two members of the PPIase family are functionally linked in yeast cells. It was found that although these proteins do not bind or catalyze the same peptides, they can generate conformational changes to substrates [33]. Another study has showed that a parvulin and a FKBP protein catalyze the *cis/trans* isomerization of peptide bonds in proteins with great homology [34]. Based on the above studies, we questioned whether the previously observed phenotypes of ΔFklB mutant strain could reverse upon expression of members of the PPIase family. To this end, we separately introduced and expressed plasmids that contained each gene belonging to the PPIase family; *fkpA, slyD, fkpB, tig, ppiA, ppiB, surA, ppiC* and *ppiD* into the ΔFklB mutant strain and we compared the ability of each one of swarming, swimming and biofilm formation (Fig. 4).

**Figure 4.**
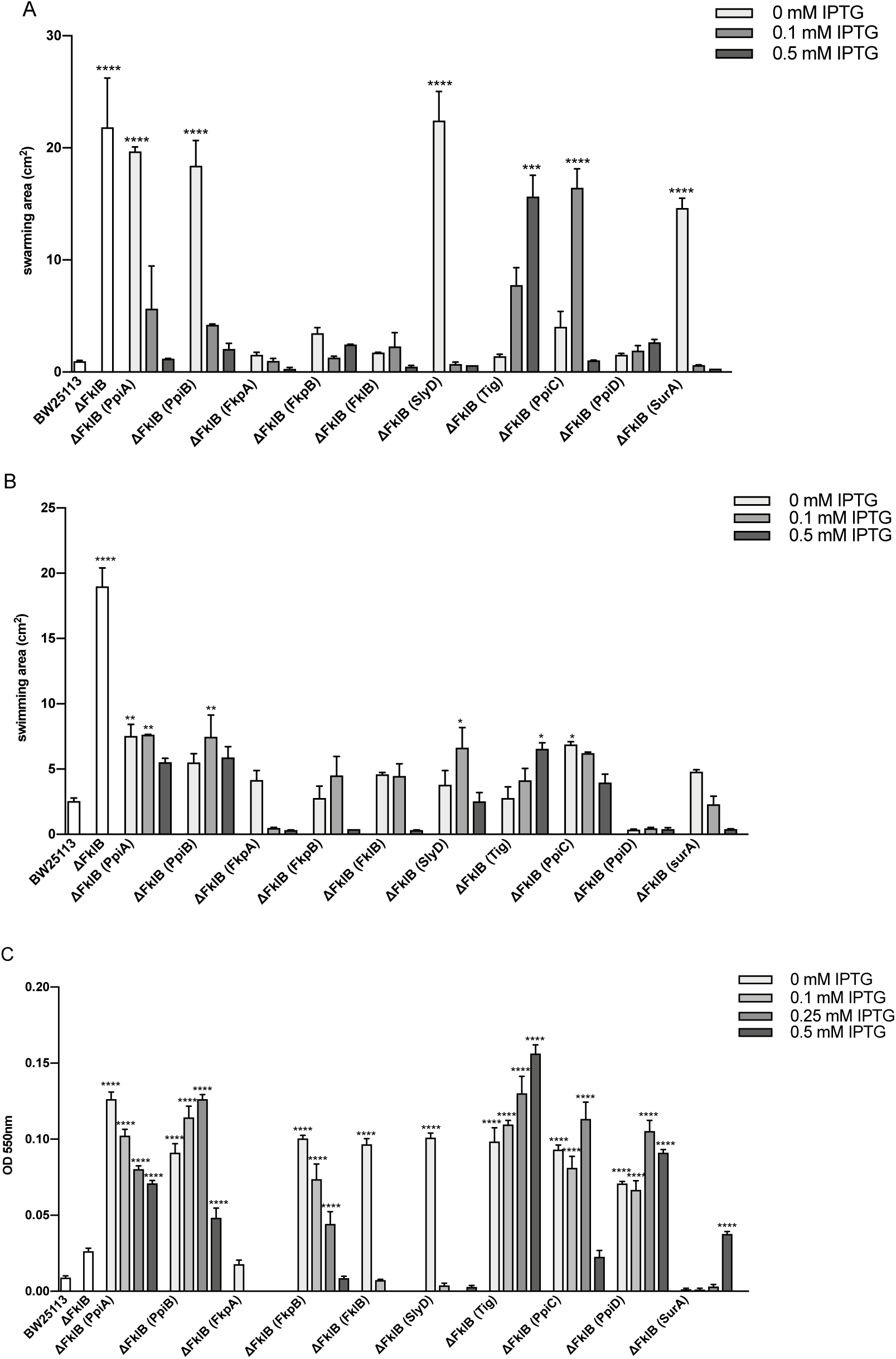
Members of the prolyl isomerase family restore the FklB mutant strain phenotypes. Swarming area (A), swimming area (B), and biofilm formation (C) of ΔFklB and ΔFklB overexpressing each PPIase family member, PpiA, PpiB, PpiC, FkpA, FkpB, FklB, SlyD, Tig, PpiC, PpiD, SurA in pCA24N vector.

Interestingly, we found that the multiple copies of all gene members of the PPIase family (0.5 mM IPTG), but not the member tig of the FKBP family, could rescue the hyper-swarming phenotype of the ΔFklB mutant (Fig 4A). The hyper-swarmer phenotype of ΔFklB was restored to wild-type levels even at single copies of genes *fkpA, ppiD*, and of course *fklB* (0 mM IPTG). This observation could be evidential of a functional overlap between FKBPs and parvulins, in swarming bacteria, perhaps hinting at the *cis/trans* isomerization of some common substrates.

The hyper-swimmer phenotype of ΔFklB was rescued upon expression of the majority of PPIases, excluding the cyclophilins ppiA and ppiB. Every member of the FKBP family was able to complement FklB’s function in swimming cells even in single copies (0 mM IPTG). There was a significant increase detected at high expression levels of tig (0.5 mM IPTG), which indicates a unique involvement of the trigger factor protein in swimming motility (Fig. 4B).

Lastly, we checked the biofilm formation phenotype of the ΔFklB could be abrogated by members of the PPIase family. We found that the expression of a great number of PPIases was not able to functionally replace FklB. The members PpiA, PpiB, FkpB, Tig, PpiC, and PpiD did not rescue the increased biofilm of the mutant strain ΔFklB, at low levels of expression (0 mM and 0.1 mM IPTG). Wild-type biofilm levels were recovered in the high-copy presence of PpiB, PpiC, and SurA (0.5 mM IPTG) and in the intermediate-copy presence of FklB and FkpB (0.25 mM IPTG). Interestingly, we identified a biofilm suppression phenotype after expressing high-levels of FKBP encoding genes, *fkpA, fkpB, slyD*, and *fklB* (0.5 mM IPTG) (Fig. 4C).

The evidence from the above experiments point towards the idea that members of the PPIase family can compensate for the absence of FklB, in swarmer, swimmer and biofilm *E. coli* cells. This functional replacement is even possible at very low copy numbers of PPIases, suggesting that there might be a substrate regulation pathway shared within the PPIase family. The data also indicate that there is a stronger physiological function commonality among members of the FKBP family through the regulation of cellular processes, post-translationally [26].

### FklB expression causes cell morphology alterations

Swarmer cells are described as elongated and hyperflagellated cells, that are able to migrate towards the edge of a swarming plate or a nutrient-rich surface, away from the initial colony [35] [36] [37]. We have previously examined E. coli cells in a planktonic phase that lack or overexpress the cyclophilin PpiB and found that in both cases the present an impaired cell division [38].

In this study, we examined the cell morphology of the control BW25113 and of the mutant ΔFklB, as well as of the strains that overexpress FklB; BW25113(FklB), BW25113(Y181A), ΔFklB(FklB) and ΔFklB(Y181A), during swarming and swimming motility. The expression of plasmids that carried the *fklB* gene and its mutant, *Y181A*, was performed in the presence of 0.1, 0.25, and 0.5 mM IPTG. We microscopically observed all the above strains after Gram staining and after DAPI staining, using an optical and a fluorescence microscope, respectively, showing the expression of FklB and Y181A at 0.25 mM IPTG (Fig. 5, 6).

**Figure 5.**
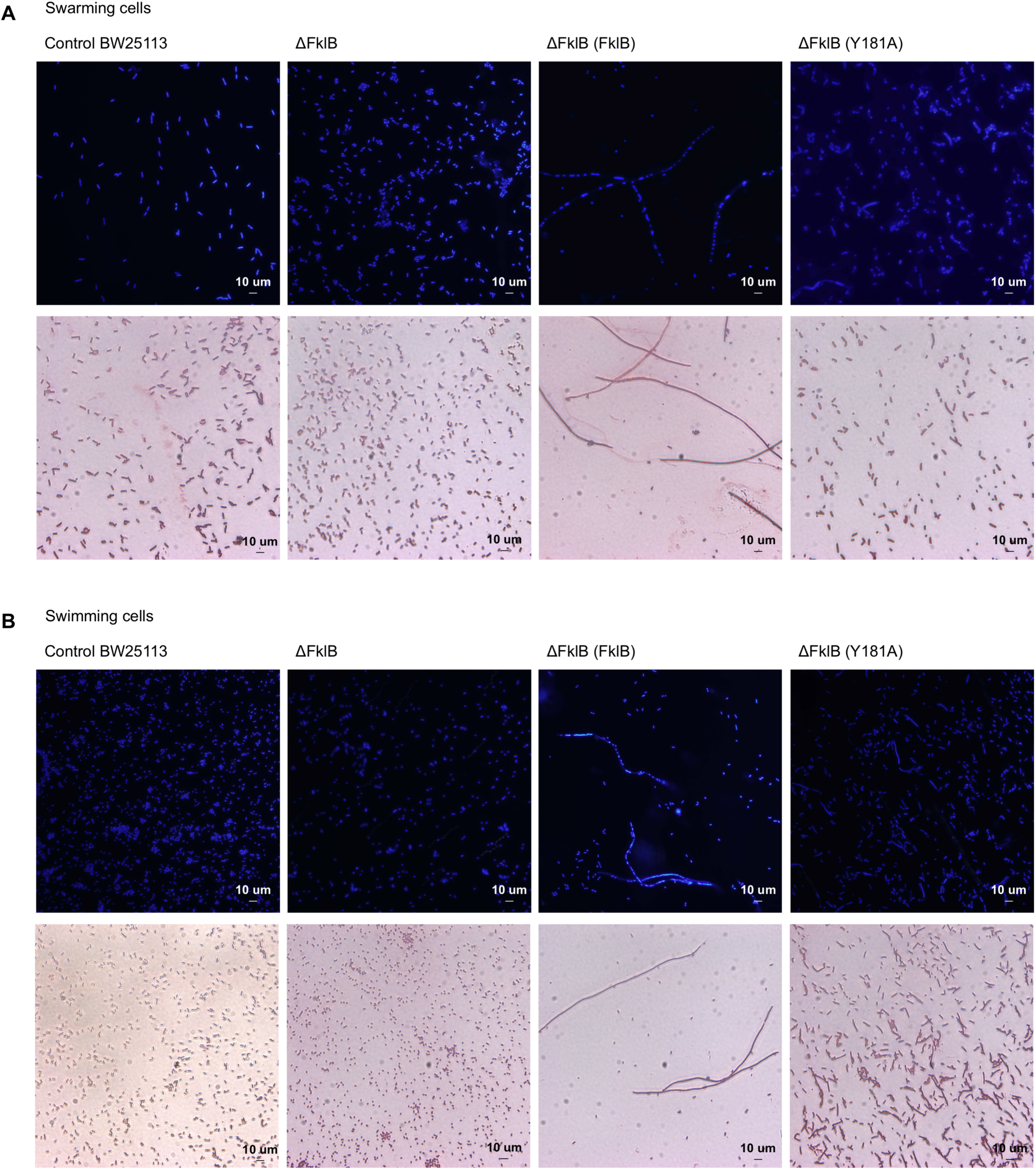
Overexpression of FklB, but not Y181A, in ΔFklB causes a cell elongation phenotype in swarming and swimming *E. coli*. ΔFklB and ΔFklB overexpressing FklB or Y181A in pCA24N vector taken from swarming (A) and swimming (B) cells were examined after DAPI (upper row) or Gram (bottom row) staining by light and fluorescent microscopy and compared to the control, BW25113. Bars represent 10 um.

**Figure 6.**
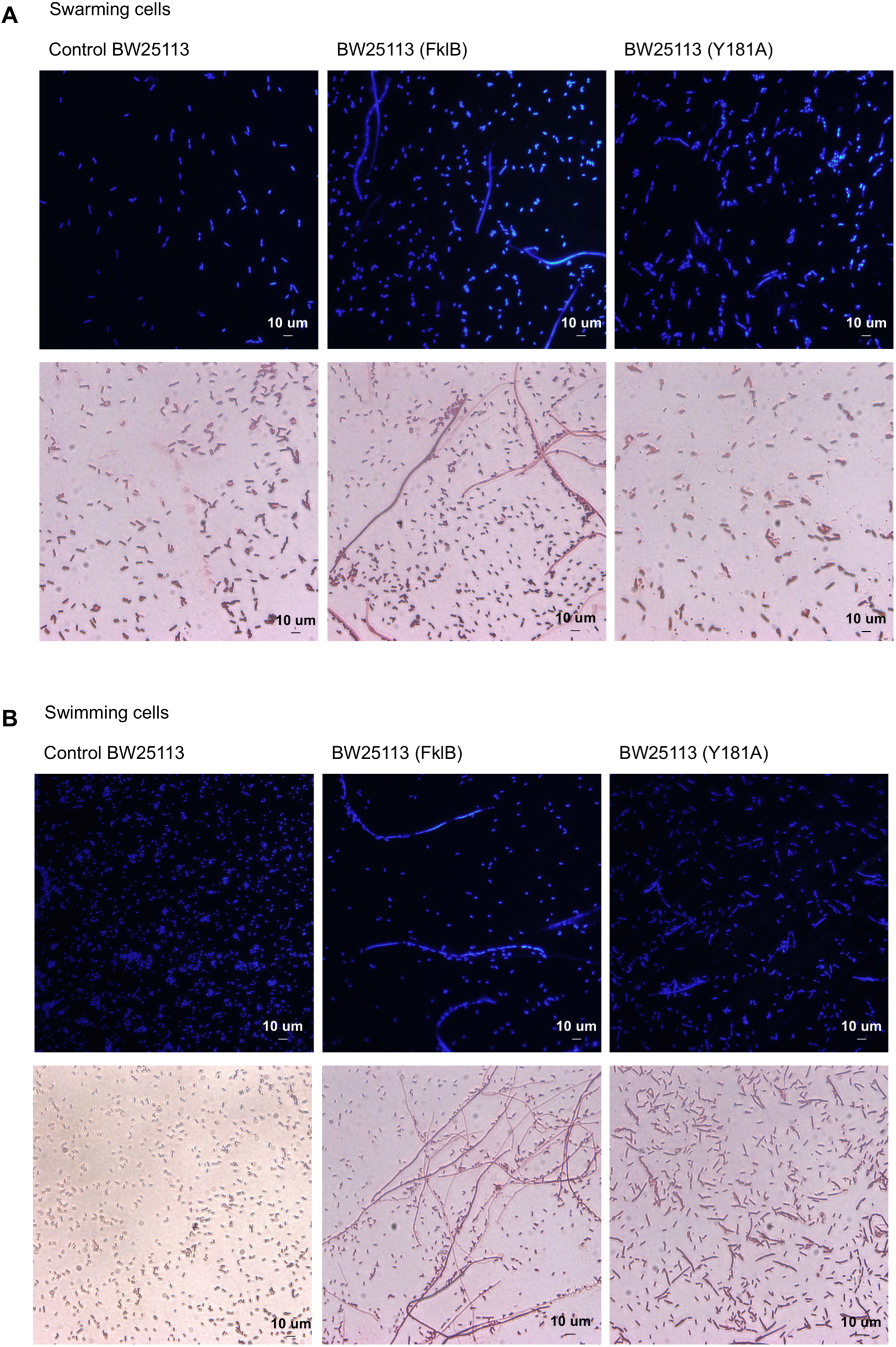
Overexpression of FklB, but not Y181A, in BW25113 causes a cell elongation phenotype in swarming and swimming *E. coli*. BW25113 overexpressing FklB or Y181A in pCA24N vector taken from swarming (A) and swimming (B) cells were examined after DAPI (upper row) or Gram (bottom row) staining by light and fluorescent microscopy and compared to the control, BW25113. Bars represent 10 um.

In swarming cells, we observed that the absence of the *fklB* gene (strain ΔFklB) did not result in a differentiated cell phenotype when compared to the control strain (Fig. 5A). However, overexpression of the *fklB* gene, in both the wild-type and mutant strains, BW25113(FklB) and ΔFklB(FklB), resulted into a phenotype characterized by elongated cells that have stopped dividing (Fig. 5A, 6A).

Additionally, a mixed population of normal size cells and cells that were not dividing was noted in the swarmer cells of strains BW25113(FklB) and ΔFklB(FklB) overexpressing the *fklB* gene (Fig. 5A, 6A). For these cells, we noticed an abnormality in the septa formation, that did not allow the separation of the cellular membrane for cell division.

Regarding the swimming motility, we noted that the phenotype of the mutant strain ΔFklB did not differ profoundly from the wild-type strain, BW25113. However, overexpression of the *fklB* gene in both strains, BW25113(FklB) and ΔFklB(FklB), caused a pronounced cell elongation, in which it appeared that the cell division had been inhibited. Figure 5 shows that the cells of strain ΔFklB(FklB) formed multiple nucleoids, therefore we concluded that the replication of the genetic material was being done normally, while the cell wall and plasma membrane separation were not permitted (Fig. 5B, 6B).

The phenotype in both swarming and swimming cells seemed to reverse after the expression of the mutant gene, *Y181A*. Strains BW25113(Y181A) and ΔFklB(Y181A) had a normal cell appearance under swarming and swimming conditions, that phenotypically corresponded to the wild-type strain. This observation led us to the conclusion that the cellular elongation of the strains BW25113(FklB) and ΔFklB(FklB) was due to the increased levels of the FklB protein. Summing up the results, it was concluded that the accumulation of the FklB protein, in the swarmer and swimmer cells, caused phenotype alterations that were specific to the increased PPIase activity (Fig. 5, 6).

## Materials and Methods

### *E. coli* strains, growth conditions and growth rate assay

The bacterial strains and plasmids that were used in this study are described in Table 1. *E. coli* K-12 BW25113 and single-gene knockout mutants [39] were obtained from the *E. coli* Genetic Stock Center. Plasmid pCA24N, as well as plasmids pCA24N containing the PPIase encoding genes were obtained from the ASKA library of the NARA Institute [40]. Unless stated otherwise, bacteria were cultivated routinely in LB (Luria–Bertani) agar or broth at 37°C with aeration. When necessary, media were supplemented with chloramphenicol (25 μg/ml) or kanamycin (25 μg/ml) or ampicillin (100 μg/ml). The specific growth rates of the *E. coli* wild-type and mutant strains were determined by measuring the turbidity at 600 nm and C.F.U/ml for two independent cultures of each strain as a function of time with turbidity values less than 0.9.

### Plasmids

The coding sequence of *Ec*FklB (NC_000913.3) was amplified using PCR and *E. coli* genomic DNA as a template. The primers used are *Ec*.b4207.H.F: 5’-CCAGGATCCGACCACCCCAACTTTTGACACC −3’ and *Ec*.b3349.H.R: 5’-CGCAAGCTTTTAGAGGATTTCCAGCAGTTC −3’. The fragment excised from amplified *Ec*FklB sequence was cloned between BamHI and HindIII sites of pPROEX-HTa, resulting in pPROEX-HTa FklB. Y181A point mutation in FklB was engineered using the gene-SOE method described by Horton [41]. The mutagenic primers used are *Ec*.b4207.Y181A.F: 5’-CCGCAGGAACTGGCAGCTGGCGAGCGCGGCGCA-3’ and *Ec*.b4207.Y181A.R: 5’- TGCGCCGCGCTCGCCATATGCCAGTTCCTGCGG-3’. The primary PCR products were purified and then used as templates for the second round of PCR. The final product was introduced into pPROEX-HTa, resulting in pPROEX-HTa Y181A. The nucleotide sequence of the gene encoding the mutant protein was confirmed by Sanger sequencing.

### Heterologous expression of FklB in *E. coli* and purification of recombinant protein

*E. coli* BL21(DE3)[F– ompT gal dcm lon hsdS_B_(r_B_^-^m_B_^-^) λ(DE3 [lacI lacUV5-T7 gene 1 ind1 sam7 nin5] (Novagen, Madison, WI, USA) was used as a host strain for overproduction of a His-tagged form of FklB or Y181A. Synthesis of recombinant proteins in *E. coli* BL21 (DE3) cells was initiated by addition of 0.25 mM isopropyl 1-thio-β-D-galactopyranoside (IPTG) when the culture reached OD600 of 0.6 and continued cultivation for additional 4h at 30°C. Recombinant proteins were purified with Ni-NTA chromatography (Ni^2+^-nitrilotriacetate, Qiagen) according to the manufacturer’s instructions. To remove any imidazole and salts in the collected fractions, fractions were pooled accordingly and dialyzed against 35 mM Hepes buffer pH8.0 and 70 mM NaCl, for 12 h. Production levels and purity of the recombinant proteins were analyzed by 15% SDS-PAGE electrophoresis.

### Motility assays

Overnight cultures of the different strains were grown, standardized to an OD600 of 1.2, and 3 μl used to stab or spotted at the center of swimming and swarming plates, respectively. The swimming plates were prepared with 0.3% Fluka agar, 1% Bacto-tryptone, 0.5% Yeast Extract and 1% NaCl. The swarming motility plates were prepared with 0.5% Fluka agar, 1% Bacto-tryptone, 0.5% Yeast Extract, 1% NaCl and 0.5% glucose. When necessary, media were supplemented with chloramphenicol (25 μg/ml) or kanamycin (25 μg/ml) and the appropriate amounts of IPTG. The plates were dried for 1-2 h at room temperature before being inoculated and were scanned after 20 h incubation at 30 °C. Petri dishes were scanned and the swarming and swimming areas were measured with the imaging software ImageJ. The experiments were carried out in three replicates.

### Biofilm formation assay

The crystal violet biofilm assays were performed as previously described [4]. Briefly, BW25113 and the *fklB* mutant strains containing either pPROEX-HTa FklB or pPROEX-HTa Y181A were grown overnight in LB medium. The overnight cultures were 1:10 diluted in 100 μl of LB medium supplemented, when necessary, with appropriate concentrations of antibiotics and IPTG, and the biofilm was formed in covered 96-well microtiter dish for 20 h without shaking at 30°C. The cell suspensions were removed and turbidity was measured at OD600. The plates were washed once with sterile distilled H_2_O to remove unbound bacteria and stained with 200 μl crystal violet (0.1% solution) for 20 min. Quantification was conducted by suspending the crystal violet stained cells in 200 μl of 20% acetone (in ethanol). Total biofilm formation was normalized by cell growth (turbidity at 600 nm) to avoid overestimating changes due to growth effects. As controls, BW25113 *fklB* mutants with empty or pPROEX-HTa or pCA24N were used.

### Peptidyl-prolyl *cis/trans* isomerase enzymatic assay

PPIase activity was tested using a chymotrypsin-coupled PPIase assay [19]. In this assay we measured the ability of FklB or Y181A to convert the *cis* isomer of the synthetic oligopeptide substrate *N*-Suc-Ala-Leu-Pro-Phe-*p*-nitroanilide into the *trans* form. The assay reaction contained 50 mM Hepes buffer pH 8.0 and 100 mM NaCl, 50 μg α-chymotrypsin (dissolved in 1 mM HCl) (Fluka), 25 μM Suc-AAPF-pNA (5 mM stock dissolved in trifluoroethanol supplemented with 0.45 M LiCl) and the appropriate amount of recombinant FklB or Y181A. The reaction was monitored at 4°C by the increase in absorbance at 390 nm (corresponding to the release of *p*-nitroanilide) using a HITACHI U-2800 spectrophotometer.

### Citrate synthase thermal aggregation assay

Thermal denaturation of citrate synthase (0.25 μM final concentration, Sigma) was achieved by incubation at 45°C, in 40 mM Hepes pH: 7.5, for 15-20 min, in the absence or in the presence of additional proteins, as previously described [21]. Aggregation of citrate synthase was measured by monitoring the increase in turbidity at 500 nm in a HITACHI U-2800 spectrophotometer equipped with a thermostatic cell holder. The absorbance change recorded is due to the increase in light scattering upon aggregation of citrate synthase. Protein disulfide isomerase (Sigma) was used in positive control reactions and albumin (Research Organics) was used in negative control reactions.

## References

[1] N. Verstraeten et al., “Living on a surface: swarming and biofilm formation,” Trends in Microbiology, vol. 16, no. 10, pp. 496–506, Oct. 2008.

[2] D. B. Kearns, “A field guide to bacterial swarming motility,” Nat. Rev. Microbiol., vol. 8, no. 9, pp. 634–644, Sep. 2010.

[3] G. L. Hazelbauer, J. J. Falke, and J. S. Parkinson, “Bacterial chemoreceptors: high-performance signaling in networked arrays,” Trends Biochem. Sci., vol. 33, no. 1, pp. 9–19, Jan. 2008.

[4] G. A. O’Toole and R. Kolter, “Flagellar and twitching motility are necessary for Pseudomonas aeruginosa biofilm development,” Mol. Microbiol., vol. 30, no. 2, pp. 295–304, Oct. 1998.

[5] M. E. Davey and G. A. O’toole, “Microbial biofilms: from ecology to molecular genetics,” Microbiol. Mol. Biol. Rev., vol. 64, no. 4, pp. 847–867, Dec. 2000.

[6] S. W. Singer et al., “Posttranslational modification and sequence variation of redox-active proteins correlate with biofilm life cycle in natural microbial communities,” ISME J, vol. 4, no. 11, pp. 1398–1409, Nov. 2010.

[7] Z. Li et al., “Diverse and divergent protein post-translational modifications in two growth stages of a natural microbial community,” Nat Commun, vol. 5, p. 4405, Jul. 2014.

[8] S. L. Rutherford and C. S. Zuker, “Protein folding and the regulation of signaling pathways,” Cell, vol. 79, no. 7, pp. 1129–1132, Dec. 1994.

[9] C. B. Kang, Y. Hong, S. Dhe-Paganon, and H. S. Yoon, “FKBP family proteins: immunophilins with versatile biological functions,” Neurosignals, vol. 16, no. 4, pp. 318–325, 2008.

[10] P. Vittorioso, R. Cowling, J. D. Faure, M. Caboche, and C. Bellini, “Mutation in the Arabidopsis PASTICCINO1 gene, which encodes a new FK506-binding protein-like protein, has a dramatic effect on plant development,” Mol. Cell. Biol., vol. 18, no. 5, pp. 3034–3043, May 1998.

[11] A. Breiman and I. Camus, “The involvement of mammalian and plant FK506-binding proteins (FKBPs) in development,” Transgenic Res., vol. 11, no. 4, pp. 321–335, Aug. 2002.

[12] M. Geisler and A. Bailly, “Tête-à-tête: the function of FKBPs in plant development,” Trends Plant Sci., vol. 12, no. 10, pp. 465–473, Oct. 2007.

[13] M. Gaestel, “Molecular chaperones in signal transduction,” Handb Exp Pharmacol, no. 172, pp. 93–109, 2006.

[14] J. U. Rahfeld et al., “Isolation and amino acid sequence of a new 22-kDa FKBP-like peptidyl-prolyl cis/trans-isomerase of Escherichia coli. Similarity to Mip-like proteins of pathogenic bacteria,” J. Biol. Chem., vol. 271, no. 36, pp. 22130–22138, Sep. 1996.

[15] N. Zang et al., “Requirement of a mip-like gene for virulence in the phytopathogenic bacterium Xanthomonas campestris pv. campestris,” Mol. Plant Microbe Interact., vol. 20, no. 1, pp. 21–30, Jan. 2007.

[16] C. Budiman, K. Bando, C. Angkawidjaja, Y. Koga, K. Takano, and S. Kanaya, “Engineering of monomeric FK506-binding protein 22 with peptidyl prolyl cis-trans isomerase. Importance of a V-shaped dimeric structure for binding to protein substrate,” FEBS J., vol. 276, no. 15, pp. 4091–4101, Aug. 2009.

[17] B. Jana, A. Bandhu, R. Mondal, A. Biswas, K. Sau, and S. Sau, “Domain structure and denaturation of a dimeric Mip-like peptidyl-prolyl cis-trans isomerase from Escherichia coli,” Biochemistry, vol. 51, no. 6, pp. 1223–1237, Feb. 2012.

[18] C. Budiman, C. Angkawidjaja, H. Motoike, Y. Koga, K. Takano, and S. Kanaya, “Crystal structure of N-domain of FKBP22 from Shewanella sp. SIB1: dimer dissociation by disruption of Val-Leu knot,” Protein Sci., vol. 20, no. 10, pp. 1755–1764, Oct. 2011.

[19] J. L. Kofron, P. Kuzmic, V. Kishore, E. Colón-Bonilla, and D. H. Rich, “Determination of kinetic constants for peptidyl prolyl cis-trans isomerases by an improved spectrophotometric assay,” Biochemistry, vol. 30, no. 25, pp. 6127–6134, Jun. 1991.

[20] C. Budiman, T. Tadokoro, C. Angkawidjaja, Y. Koga, and S. Kanaya, “Role of polar and nonpolar residues at the active site for PPIase activity of FKBP22 from Shewanella sp. SIB1,” FEBS J., vol. 279, no. 6, pp. 976–986, Mar. 2012.

[21] J. Buchner, H. Grallert, and U. Jakob, “Analysis of chaperone function using citrate synthase as nonnative substrate protein,” Meth. Enzymol., vol. 290, pp. 323–338, 1998.

[22] R. P. Jakob, G. Zoldák, T. Aumüller, and F. X. Schmid, “Chaperone domains convert prolyl isomerases into generic catalysts of protein folding,” Proc. Natl. Acad. Sci. U.S.A., vol. 106, no. 48, pp. 20282–20287, Dec. 2009.

[23] G. Zoldák and F. X. Schmid, “Cooperation of the prolyl isomerase and chaperone activities of the protein folding catalyst SlyD,” J. Mol. Biol., vol. 406, no. 1, pp. 176–194, Feb. 2011.

[24] A. Galat, “Peptidylprolyl cis/trans isomerases (immunophilins): biological diversity--targets--functions,” Curr Top Med Chem, vol. 3, no. 12, pp. 1315–1347, 2003.

[25] T. Inoue, R. Shingaki, S. Hirose, K. Waki, H. Mori, and K. Fukui, “Genome-wide screening of genes required for swarming motility in Escherichia coli K-12,” J. Bacteriol., vol. 189, no. 3, pp. 950–957, Feb. 2007.

[26] A. Skagia, C. Zografou, E. Vezyri, A. Venieraki, P. Katinakis, and M. Dimou, “Cyclophilin PpiB is involved in motility and biofilm formation via its functional association with certain proteins,” Genes Cells, vol. 21, no. 8, pp. 833–851, Aug. 2016.

[27] A. Skagia et al., “Structural and functional analysis of cyclophilin PpiB mutants supports an in vivo function not limited to prolyl isomerization activity,” Genes Cells, vol. 22, no. 1, pp. 32–44, Jan. 2017.

[28] M. M. Pearson, D. A. Rasko, S. N. Smith, and H. L. T. Mobley, “Transcriptome of swarming Proteus mirabilis,” Infect. Immun., vol. 78, no. 6, pp. 2834–2845, Jun. 2010.

[29] M. J. McBride and T. F. Braun, “GldI is a lipoprotein that is required for Flavobacterium johnsoniae gliding motility and chitin utilization,” J. Bacteriol., vol. 186, no. 8, pp. 2295–2302, Apr. 2004.

[30] M. Klausen et al., “Biofilm formation by Pseudomonas aeruginosa wild type, flagella and type IV pili mutants,” Mol. Microbiol., vol. 48, no. 6, pp. 1511–1524, Jun. 2003.

[31] N. C. Caiazza, R. M. Q. Shanks, and G. A. O’Toole, “Rhamnolipids modulate swarming motility patterns of Pseudomonas aeruginosa,” J. Bacteriol., vol. 187, no. 21, pp. 7351–7361, Nov. 2005.

[32] A. Skagia et al., “Structure-Function Analysis of the Periplasmic Escherichia coli Cyclophilin PpiA in Relation to Biofilm Formation,” J. Mol. Microbiol. Biotechnol., vol. 27, no. 4, pp. 228–236, 2017.

[33] F. Fujimori et al., “Crosstalk of prolyl isomerases, Pin1/Ess1, and cyclophilin A,” Biochem. Biophys. Res. Commun., vol. 289, no. 1, pp. 181–190, Nov. 2001.

[34] Y. Mamane, S. Sharma, L. Petropoulos, R. Lin, and J. Hiscott, “Posttranslational regulation of IRF-4 activity by the immunophilin FKBP52,” Immunity, vol. 12, no. 2, pp. 129–140, Feb. 2000.

[35] C. Allison and C. Hughes, “Bacterial swarming: an example of prokaryotic differentiation and multicellular behaviour,” Sci Prog, vol. 75, no. 298 Pt 3–4, pp. 403–422, 1991.

[36] R. M. Harshey, “Bees aren’t the only ones: swarming in gram-negative bacteria,” Mol. Microbiol., vol. 13, no. 3, pp. 389–394, Aug. 1994.

[37] L. McCarter and M. Silverman, “Surface-induced swarmer cell differentiation of Vibrio parahaemolyticus,” Mol. Microbiol., vol. 4, no. 7, pp. 1057–1062, Jul. 1990.

[38] A. Skagia, C. Zografou, A. Venieraki, C. Fasseas, P. Katinakis, and M. Dimou, “Functional analysis of the cyclophilin PpiB role in bacterial cell division,” Genes Cells, vol. 22, no. 9, pp. 810–824, Sep. 2017.

[39] T. Baba et al., “Construction of Escherichia coli K-12 in-frame, single-gene knockout mutants: the Keio collection,” Mol. Syst. Biol., vol. 2, p. 2006.0008, 2006.

[40] M. Kitagawa et al., “Complete set of ORF clones of Escherichia coli ASKA library (a complete set of E. coli K-12 ORF archive): unique resources for biological research,” DNA Res., vol. 12, no. 5, pp. 291–299, 2005.

[41] R. M. Horton, H. D. Hunt, S. N. Ho, J. K. Pullen, and L. R. Pease, “Engineering hybrid genes without the use of restriction enzymes: gene splicing by overlap extension,” Gene, vol. 77, no. 1, pp. 61–68, Apr. 1989.

